# *coronapp*: A Web Application to Annotate and Monitor SARS-CoV-2 Mutations

**DOI:** 10.1101/2020.05.31.124966

**Authors:** Daniele Mercatelli, Luca Triboli, Eleonora Fornasari, Forest Ray, Federico M. Giorgi

**Author notes:** Corresponding author. (Giorgi FM).

## Abstract

The avalanche of genomic data generated from the SARS-CoV-2 virus requires the development of tools to detect and monitor its mutations across the world. Here, we present a webtool, *coronapp*, dedicated to easily processing user-provided SARS-CoV-2 genomic sequences and visualizing current worldwide status of SARS-CoV-2 mutations.

The webtool allows users to highlight mutations and categorize them by frequency, country, genomic location and effect on protein sequences, and to monitor their presence in the population over time.

The tool is available at http://giorgilab.unibo.it/coronapp/ for the worldwide dataset and at http://giorgilab.unibo.it/coronannotator/ for the annotation of user-provided sequences. The full code is freely shared at https://github.com/federicogiorgi/giorgilab/tree/master/coronapp

**Data Availability Statement:** The data that support the findings of this study derive from the GISAID consortium and are openly available in Github, in Rdata format for the R environment, in files results.rda and metadata.rda, at the following link: https://github.com/federicogiorgi/giorgilab/tree/master/coronapp/data

## Introduction

SARS-CoV-2 is a novel pathogenic enveloped RNA beta-coronavirus causing a severe illness in human hosts known as coronavirus disease-2019 (COVID-19). The predominant COVID-19 illness is a viral pneumonia, often requiring hospitalization and in some cases intensive care [1]. With almost 27.5 million laboratory-confirmed positive cases worldwide as of 9 September 2020 and an estimated case fatality rate across 204 countries of 5.2%, COVID-19 has become a global health challenge in only a few months [2]. SARS-CoV-2 infection depends on the recognition of host angiotensin converting enzyme 2 (ACE2), exposed on the cell surface in human lung tissues [3,4]. SARS-CoV-2 spike glycoprotein binds ACE2, mediating membrane fusion and cell entry [5]. Upon cell entry, the virus subverts host cell molecular processes, inducing interferon responses and eventually apoptosis [6].

To date, much effort has been made to develop therapeutic strategies to limit SARS-CoV-2 transmission and replication, but no treatment or vaccine has proven effective against the virus, and repurposing of approved therapeutic agents has been the main practical approach to manage the emergency so far [7]. As viruses mutate during replication, the emergence of SARS-CoV-2 sub-strains and the challenge of a probable antigenic drift require attention, especially for vaccine development [8].

Although sequence analyses of SARS-CoV-2 have shown that genomic variability is very low [9], new SARS-CoV-2 mutation hotspots are emerging due to the high number of infected individuals across countries and to viral replication rates [10]. Three major SARS-CoV-2 clades known as clade G, V, and S have emerged, showing a different geographical prevalence [10]. The most frequent mutation detected so far defines the G clade and causes an aminoacidic change, aspartate (D) or glycine (G), at position 614 (D614G) of the viral Spike protein [11].

Continual genomic surveillance should be considered to monitor the possible appearance of viral subtypes characterized by altered tropism, or causing more aggressive symptoms. Constant and widespread monitoring of mutations is also a powerful means of informing drug development and global or local pandemic management. The Global Initiative on Sharing All Influenza Data (GISAID) has collected to date (9 September 2020) over 90,000 publicly accessible SARS-CoV-2 sequences. The GISAID effort has made it possible to compare genomes on a geographical and temporal scale and an increasing number of laboratories have started to sequence COVID-19 patient samples worldwide [12,13]. Several online tools have been developed to monitor the evolution of the virus from a phylogenetic perspective, such as Nextstrain [14], or to visualize epidemiological data such as number of cases and deaths [15]. However, no online tool currently exists to annotate user-provided SARS-CoV-2 genomic sequences, which may derive from specific GISAID subsets or from sequencing efforts of individual laboratories. Neither does any tool specifically monitor the prevalence of specific SARS-CoV-2 mutations associated to particular geographic regions or protein locations, nor their frequency in the population over time.

To overcome these limitations, we have developed *coronapp*, a web application with two purposes: real-time tracking of SARS-CoV-2 mutational status and annotation of user-provided viral genomic sequences. Our tool enables users to easily perform genomic comparisons and provides an instrument to monitor SARS-CoV-2 genomic variance, both worldwide and by uploading custom and locally produced genomic sequences. The webtool is available at http://giorgilab.dyndns.org/coronapp/ and the full source code is shared on Github https://github.com/federicogiorgi/giorgilab/tree/master/coronapp

## Results

The webtool *coronapp* is available at the website http://giorgilab.dyndns.org/coronapp/ and it automatically provides the user with the current status of SARS-CoV-2 mutations worldwide. The app also allows users to annotate user-provided sequences (Figure 1 A). There are multiple functionalities of *coronapp*, described in the following paragraphs.

**Figure 1.**
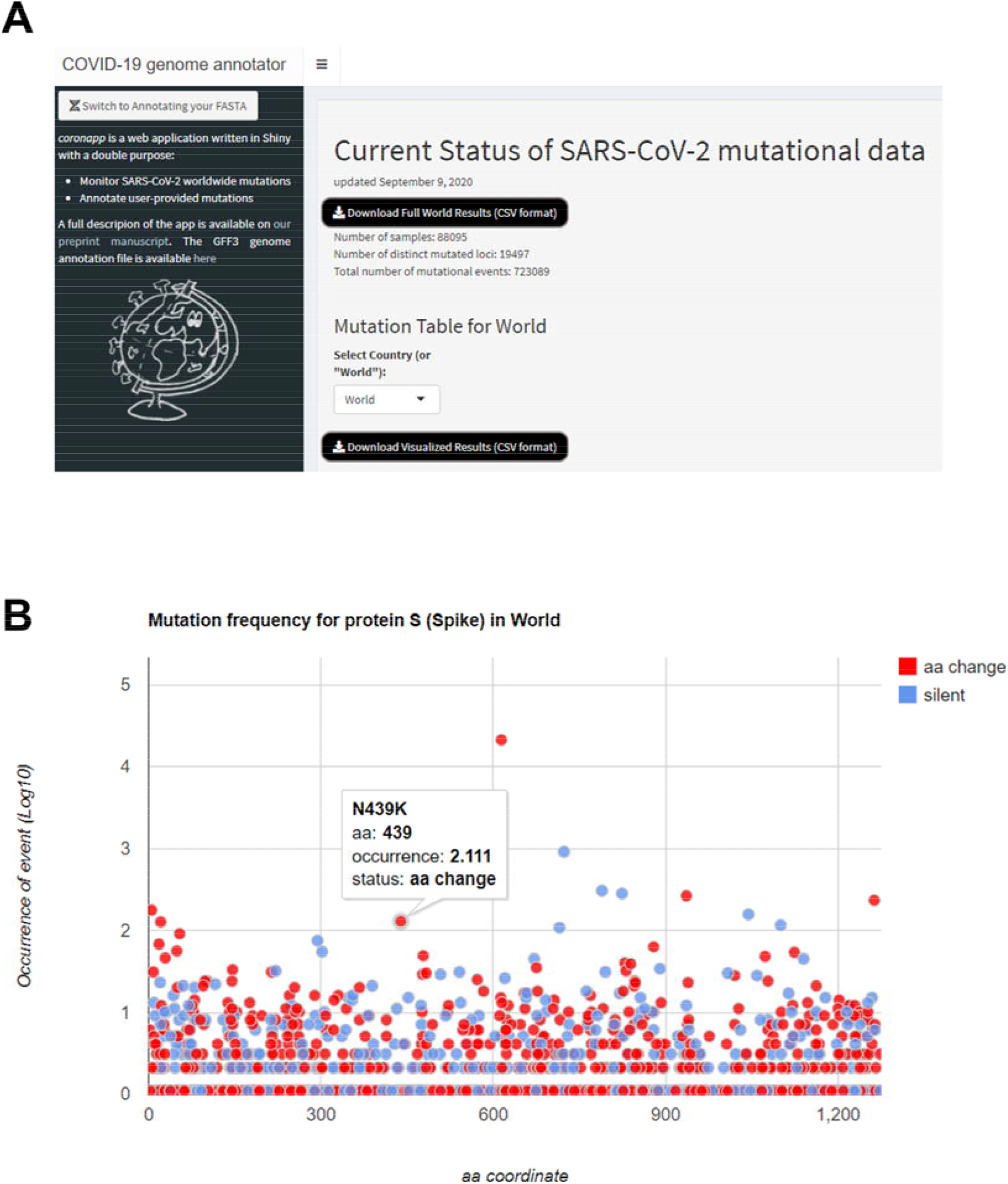
Overview of *coronapp*. **A**. Screenshot of the entry page of *coronapp* showing the basic tool description, the interface to upload user-provided sequences and the overall summary of the mutations detected worldwide. **B**. Common interface showing mutation frequency in SARS-CoV-2 proteins, with occurrence of the mutation on the Y-axis and protein coordinate on the Y-axis. Red dots indicate amino acid (aa)-changing mutations, and blue dots indicate silent mutations. Tooltip functionality is also provided to identify and quantify each mutation on mouse-over.

### Current Status of SARS-CoV-2 mutational data

A worldwide analysis is shown, generated using data from GISAID. Specifically, we processed all SARS-CoV-2 complete (>29,000 sequenced nucleotides) genomic sequences, excluding low-quality sequences (>5% undefined nucleotide “N”) and viruses extracted from non-human hosts.

The underlying database is updated weekly, and we provide the date of the last version as a reference for studies based on the data provided. We indicate the number of samples processed and the total number of mutational events detected (Figure 1 A). We also show the number of distinct mutated loci. Currently, this number is slightly below 20,000, meaning that two thirds of the original Wuhan SARS-CoV-2 genome has been affected by mutations and/or sequencing errors (the full length of the reference genome is 29,903 nucleotides, based on sequence id NC_045512.2).

### Mutation frequency in SARS-CoV-2 proteins

We show the frequency of mutations along the length of every SARS-CoV-2 protein, reporting in the X-axis the amino acid position and on the Y-axis its frequency, either as number of observed samples carrying the mutation, the vase 10 logarithm of that number, or the percentage over all sequenced samples. In the example in Figure 1 B, we show the most frequent mutations affecting the viral Spike protein S, distinguishing silent mutations and amino acid-changing mutations (including the introduction of STOP codons). For Spike, the mutations appear to be evenly distributed in frequency along the protein length, with the most frequent mutation being the aforementioned D614G. Mouse-over functionality is provided to allow the user to identify the selected mutation (e.g. N439K in Figure 1 B).

### The SARS-CoV-2 mutation table

The user can visualize or download the full table of mutations on which the webtool operates (Figure 2 A). This table is frequently updated and allows the user to specify a worldwide or a country-specific dataset. The table also provides a Search function to look for specific variants or sample ids, and it can be viewed online or downloaded in full as a Comma-Separated Values (CSV) file.

**Figure 2.**
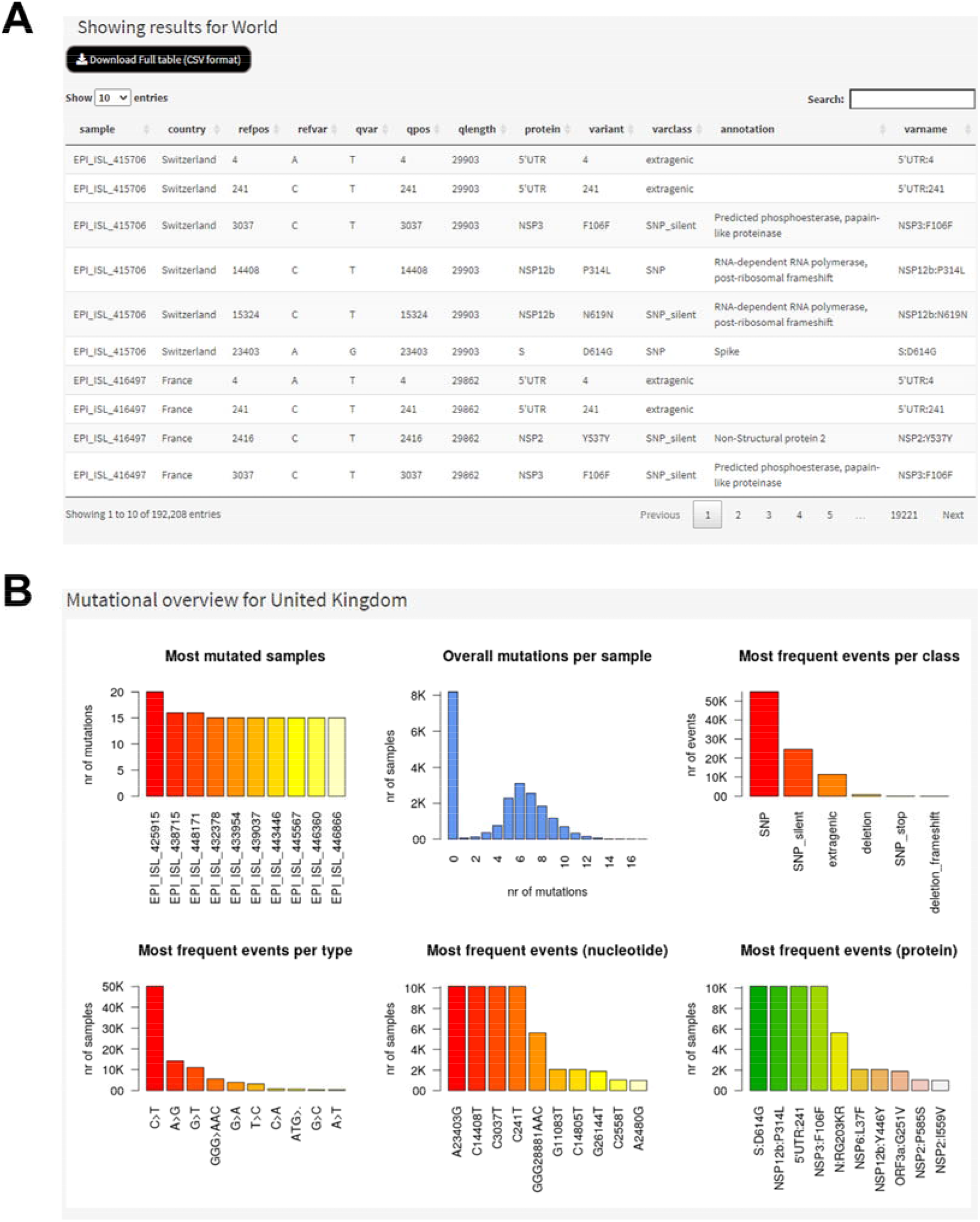
Mutation table and overview in *coronapp*. **A**. Result table of *coronapp*, available both for worldwide-precomputed and user-input analyses. A “download full table” button is provided to allow the user to perform larger-scale analyses autonomously. **B**. Barplots showing the most mutated samples, overall sample mutations and most frequent mutation events, classes and types. This analysis is also available both for worldwide-precomputed and user-input analyses.

The table shows every mutation in a specific geographical area, reporting:

- the GISAID sample ID (useful for cross-reference with the GISAID database and other analyses based on it, e.g. Nexstrain).
- The country where the sample was collected.
- The position of the mutation, on the reference genome (refpos) and on the sample (qpos).
- The sequence at the mutation site, on the reference genome (refvar) and on the sample (qvar).
- The length of the sample genome (qlength); the reference genome is 29,903 nucleotides long.
- The protein affected by the mutation or, if the mutation is extragenic, the denomination of the untranslated region (UTR), e.g. 5’UTR or 3’UTR.
- The effect of the mutation on the amino acid sequence of the protein (variant). This uses the canonical mutational standard, indicating the original amino acid(s), the position on the protein, and the mutated amino acid(s). An asterisk (*) indicates a STOP codon, while the letters indicate amino acids in IUPAC code. E.g. a mutation P315L indicates a leucine mutation (L) on the amino acid location 315, normally occupied by a proline (P). Nucleotide mutations can be silent, i.e. not yielding any aminoacidic change, e.g. the mutation F106F, where the codon of phenylalanine 106 is affected but without changing the corresponding amino acid. As in the previous column, mutations affecting UTR regions are simply reported as the location of the nucleotide affected.
- The class of the mutation, of which there are currently 10 types:

○ SNP: a change of one or more nucleotides, determining a change in amino acid sequence.
○ SNP_stop: a change of one or more nucleotides, yielding the generation of one or more STOP codons.
○ SNP_silent: a change of one or more nucleotides with no effect in protein sequence.
○ Insertion: the insertion of 3 (or multiples of 3) nucleotides, causing the addition of 1 or more amino acids to the protein sequence.
○ Insertion_stop: the insertion of 3 (or multiples of 3) nucleotides, causing the generation of a novel STOP codon.
○ Insertion_frameshift: the insertion of nucleotides not as multiples of 3, causing a frameshift mutation.
○ Deletion: the deletion of 3 (or multiples of 3) nucleotides, causing the removal of 1 or more amino acids to the protein sequence.
○ Deletion_stop: the removal of 3 (or multiples of 3) nucleotides, causing the generation of a novel STOP codon.
○ Deletion_frameshift: the deletion of nucleotides not as multiples of 3, causing a frameshift mutation.
○ Extragenic: a mutation affecting intergenic or UTR regions.
- The extended annotation of the protein region affected by the mutation (e.g. “Spike” for “S” or “Predicted phosphoesterase, papain-like proteinase” for NSP3, the Non-Structural Protein 3).
- The full name of the variant (varname), in the format proteinName:AApositionAA, to allow for unique denomination of viral proteome variants.

### Mutational overview

The user is also provided with a general overview of the mutational status of the selected country or the entire world (Figure 2 B). Six bar plots provide a summary and highlights of the dataset, specifically:

- The most mutated samples, indicating which samples (in GISAID IDs) carry the highest number of mutations
- The overall mutations per sample, indicating the distributions of mutations per sample. It has been previously reported [10] that the current mode for mutation number compared to the reference NC_045512.2 genome is 7.5.
- The most frequent events per class. Classes are the same as reported in the mutation table and are described in the previous paragraph.
- The most frequent events per type. Individual mutation types are shown as specific nucleotides events, e.g. cytosine to thymidine transitions (C>T), guanosine to thymidine transversion (G>T) or even multinucleotide mutations (e.g. GGG>AAC, observed in the Nucleocapsid protein). As reported before, nucleotide transitions seem to be the most abundant SARS-CoV-2 type of mutational event worldwide [11].
- The most frequent events, either in nucleotide coordinates or in aminoacidic coordinates. Currently, the most frequent events are four mutations affecting SARS-CoV-2 genomes belonging to clade G, which is the most sequenced worldwide and predominant in Europe. These mutations are A23403G (associated to the already mentioned D614G mutation in the Spike protein), C3037T, C14408T and C241T.

### Analysis of mutations over time

The *coronapp* webtool allows users to monitor the abundance and frequency of any SARS-CoV-2 mutation in any country specified (Figure 3 A). Both plots in this section report continuous dates on the X-axis, starting on the day of the first collected SARS-CoV-2 genome available on GISAID: December 24, 2019.

**Figure 3.**
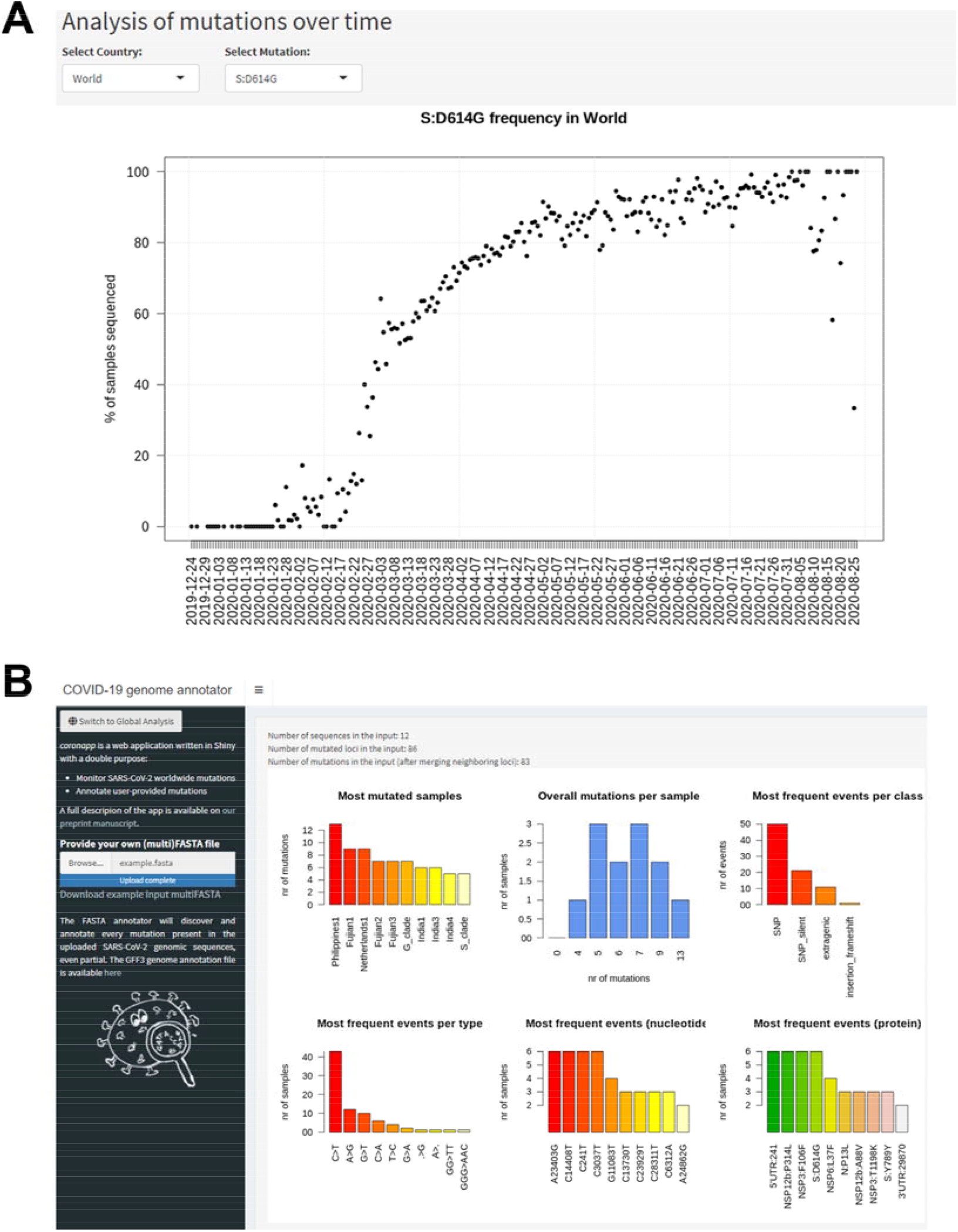
Analysis of mutations over time. **A**. Chronological analysis made by *coronapp*, showing the frequency of each user-specified mutation in any user-specified country (or worldwide). The graph shows the same data normalized by total number of samples, as the percentage of samples sequenced in a specific day and carrying the mutation. **B**. Screenshot of the *coronannotor* companion tool, allowing users to annotate their own SARS-CoV-2 sequences (provided as FASTA files).

The “abundance” plot reports on the Y-axis the number of samples carrying a selected mutation in a particular day, in the specified country or worldwide. Since the date reported is the collection date (not the submission date to the GISAID database), there is usually a drop towards the right part of the plot, as there are fewer sequences collected approaching the day of the analysis. The “frequency” plot on the other hand normalizes the abundance of mutations by the total number of sequences generated on each day. The plot currently shows a sharp increase in clade G-associated mutations (e.g. S:D614G), as these mutations are most frequent in countries where sequencing is more pervasive (e.g. United Kingdom).

### Annotation of user-provided SARS-CoV-2 genomic sequence

*coronapp* provides the user with an optional tool, *coronannotor,* providing the optional possibility of uploading one or more SARS-CoV-2 genomic sequences, which can be complete or partial. The format of the sequences is standard FASTA, and an example input FASTA containing 12 sequences is provided (Figure 3 B). The analysis is almost instantaneous and shows an overall breakdown of the most mutated samples and most frequent mutations in the dataset. Moreover, a full table of all detected mutations is provided: this can be visualized and searched on the web browser or downloaded as a standard CSV file. Finally, a mutation frequency plot is provided, allowing the user to visualize mutation frequency in selected proteins.

The user can easily return to the worldwide status of the app by refreshing or reopening the page.

## Discussion

Our webtool *coronapp* provides a fast, simple tool to annotate user-provided SARS-CoV-2 genomes and visualize all mutations currently present in viral sequences collected worldwide. The results provided by this instrument can have several applications. The main purpose of *coronapp* is to help medical laboratories at the front lines of COVID-19 fight with the opportunity to quickly define the mutational status of their sequences, even without dedicated bioinformaticians.

Additionally, it enables scientists to perform mutational co-variance analyses and to identify present and future significant functional interactions between viral mutations, as previously attempted for the influenza virus and the human immunodeficiency virus (HIV) [16]. Another application is the identification of the most frequent mutations in specific protein regions: for example, our tool can quickly identify that the most frequent mutation in the Spike protein, D614G, lies outside the known interaction domain with the human protein ACE2, which spans roughly between Spike amino acids 330 and 530 [17].

A recently published structural model simulating the effect of the D614G mutation on the 3D structure of the spike protein has suggested that this mutation may result in a viral particle which binds ACE2 receptors less efficiently, due to the masking of the host receptor binding site on viral spikes [18]. The same researchers have reported a possible correlation of the D614G form with increased case fatality rates, hypothesizing that this mutation may lead to a viral form which is better suited to escape immunologic surveillance by eliciting a lower immunologic response [18]. The *coronapp* analysis highlighted in Figure 1 B shows that a mutation located within the Spike/ACE2 interaction domain is the change of Asparagine (N) to a Lysine (K) in position 439 of the Spike sequence; this mutation could affect the protein folding or its affinity with ACE2, as Asparagine is less charged than the basic amino acid Lysine.

One of *coronapp*’s key strengths is to help prioritize scientific efforts on specific aminoacidic variations that could affect the efficacy of anti-viral strategies or the development of a vaccine by tracking the most frequent mutations in the population. A further novelty of *coronapp* is that it provides a mean to assess the growth or decline of specific mutations over time, in order to identify possible viral adaptation mechanisms.

We provide not only the webtool, but also all the underlying code for the annotation and visualization steps on a public Github repository, in order to help other computational scientists in the ongoing battle against COVID-19. Furthermore, the *coronapp* structure and concept could be expanded to other current and future pathogens as well (e.g. the seasonal influenza or HIV), in order to monitor the mutational status across proteins, countries and time.

## Materials and methods

The webtool *coronapp* has been developed using the programming language R and is based on a Shiny server (current version 1.4.0.2) running on R version 3.6.1. The app is based on two distinct files, server.R and ui.R, managing the server functionalities and the browser visualization processes, respectively. The results visualization utilizes both basic R functions and Shiny functionalities; for tooltip functionality, *coronapp* uses the R package *googleVis* v0.6.4, which provides an interface between R and the Google visualization API [19].

The core of the annotation of the user-provided sequences rests in the NUCMER (Nucleotide Mummer) alignment tool, version 3.1 [20]. Nucmer output is processed by UNIX and R scripts provided in Github within the server.R file.

## Authors’ contributions

DM drafted the manuscript and performed the mutational analysis and literature search. LT developed the user interface code and drafted the methodological parts of the manuscript. EF worked on graphical interface of the webtool. FR wrote the manuscript and performed literature search. FMG designed the study, developed the server code, finalized the manuscript and provided financial support. All authors tested the webtool and provided original contributions to its development. All authors read and approve the final manuscript.

## Acknowledgements

We thank the Italian Ministry of University and Research for their support, under the Montalcini Grant 2016.

